# Regulatory RNPs: A novel class of ribonucleoproteins that contribute to ribosome heterogeneity

**DOI:** 10.1101/128090

**Authors:** Aaron R. Poole, Ian Vicino, Hiro Adachi, Yi-Tao Yu, Michael D. Hebert

## Abstract

Ribonucleoproteins (RNPs), which are comprised of non-coding RNA and associated proteins, are involved in essential cellular processes such as translation and pre-mRNA splicing. One class of RNP is the small Cajal body-specific RNP (scaRNP), which contributes to the biogenesis of small nuclear RNPs (snRNPs) that are central components of the spliceosome. Interestingly, three scaRNAs are internally processed, generating stable nucleolus-enriched RNAs of unknown function. Here we provide evidence that these RNAs become part of novel RNPs we term regulatory RNPs (regRNPs). We postulate that regRNPs can impact rRNA modifications via interactions with the guide RNA component of small nucleolar RNPs (snoRNPs). Most modifications within rRNA (predominantly pseudouridylation and ribose 2’-*O*-methylation) are conducted by snoRNPs, and we hypothesize that the activity of at least some of these snoRNPs is under the control of regRNPs. Ribosome heterogeneity leading to specialized ribosomes is an exciting emerging concept. Because modifications within rRNA can vary in different physiological or pathological situations, rRNA modifications are thought to be the major source of ribosome heterogeneity. Our identification of regRNPs thus provides important and timely insight into how ribosome heterogeneity may be accomplished. This work also provides additional functional connections between the Cajal body and the nucleolus.

**Summary Statement:** Processed scaRNAs give rise to a novel regulatory RNP which regulates the modification of ribosomal RNA. These findings provide insight into the mechanisms governing ribosome heterogeneity.

## Introduction

Ribosomal RNA (rRNA) is extensively modified (Khan and Maden, 1978; Maden, 1972; Maden and Salim, 1974), and occurs at precise locations (Maden, 1986; Maden, 1988). Two common modifications, the majority of which are conducted by small nucleolar ribonucleoproteins (snoRNPs), are pseudouridylation and ribose 2’-*O*-methylation. There are two kinds of snoRNPs: box H/ACA (responsible for pseudouridylation) and box C/D (responsible for 2’-*O*-methylation) (Kiss, 2004). Box H/ACA or box C/D snoRNAs have an antisense region which base pairs with target rRNA and thereby facilitates the modification of rRNA target sites (Kiss, 2004). Recent evidence has shown that methylation modifications in rRNA are heterogeneous (Krogh et al., 2016) (Incarnato et al., 2017). While some sites are constitutively methylated, others may only be modified in 60% or 70% of rRNA. It is presently unknown what regulatory factors are involved in determining modification levels in rRNA. Given the vast amount of modifications present in human rRNA (approximately 100 sites of ribose methylation and 100 sites of pseudouridylation), a full characterization of the factors that orchestrate the level of rRNA modification is of fundamental importance.

A possible clue as to how the methylation of rRNA is regulated may be found in the study of small Cajal body-specific RNAs (scaRNAs). ScaRNAs were identified in the early 2000s (Darzacq et al., 2002; Tycowski et al., 2004) and localize in the Cajal body (CB), a subnuclear domain that participates in the formation of many types of RNPs (Hebert and Poole, 2016; Meier, 2016; Raimer et al., 2016; Stanek, 2016; Trinkle-Mulcahy and Sleeman, 2016). ScaRNAs are predicted to guide the modification of nucleotides in small nuclear RNA (snRNA), which are part of small nuclear RNPs (snRNPs). SnRNPs are crucial components of the spliceosome required for pre-mRNA splicing. The modifications in the snRNA component of snRNPs are essential for proper snRNP biogenesis and spliceosome function (Yu et al., 1998) (Kiss, 2004). ScaRNAs can be placed into 3 categories based on *cis*-elements in the RNA sequence and the proteins that interact with these RNAs: box C/D, box H/ACA, and mixed domain. Box C/D scaRNAs are identified by the presence of C/C’ (RUGAUGA) and D/D’ (CUGA) consensus sequences (Tycowski et al., 1996). Nop56, Nop58, 15.5 kDa, and fibrillarin, a methyltransferase, form the core protein complex on box C/D scaRNAs (Baserga et al., 1991; Fatica et al., 2000; Gautier et al., 1997; Schimmang et al., 1989; Szewczak et al., 2002; Tyc and Steitz, 1989; Watkins et al., 1996). Box H/ACA scaRNAs are defined by the presence of an H-box (ANANNA) and ACA-box (ACA) which flank a stemloop structure (Balakin et al., 1996; Bousquet-Antonelli et al., 1997; Ganot et al., 1997). Nop10, Gar1, Nhp2, and dyskerin, a pseudouridylase, form the core protein complex on box H/ACA scaRNAs (Fig. 1). Of the 28 known scaRNAs in human, three of these, all of which are box C/D, are known to be internally processed and give rise to stable, nucleolus-enriched RNA fragments: scaRNA 2, 9, and 17 (Tycowski et al., 2004). The fragments derived from these three scaRNAs are mgU2-61 (from scaRNA2), mgU2-19 and mgU2-30 (from scaRNA9) and mgU4-8 (from scaRNA17) (Fig. 1).

**Figure 1.**
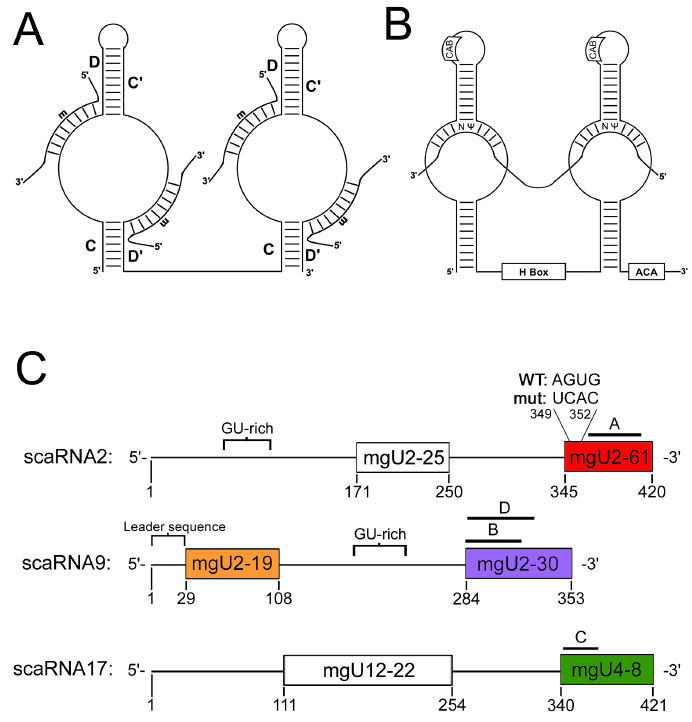
Box C/D and box H/ACA RNPs. (A) Schematic representation of box C/D scaRNA guide interaction with target RNAs promoting methylation (m). (B) Schematic representation of box H/ACA guide interaction with target RNA promoting pseudouridylation (Ψ). Note that the same arrangement between guide and target RNA is observed for box C/D and box H/ACA snoRNAs. In human, box H/ACA scaRNAs contain a *cis* element known as the CAB box which facilitates interaction of these RNAs with the protein WRAP53 and subsequent localization to CBs. (C) Schematic representation of scaRNA 2, 9, and 17. Both guide domains are shown for each scaRNA and fragments generated from each scaRNA are colored. The location of the probes used for Northern blotting are indicated by A, B, or C and the location of the probe used for RNA FISH to detect scaRNA9 is indicated by D. The GU-rich repeat region in scaRNA 2 and 9 is indicated. Also shown is the leader sequence at the 5’ end of scaRNA9 and the AGUG sequence at the beginning of the scaRNA2 mgU2-61 domain. This AGUG sequence was mutated to UCAC.

Despite numerous other proteins and processes that have been shown to be mandatory for CB integrity and composition (Lemm et al., 2006; Mahmoudi et al., 2010; Shpargel and Matera, 2005; Zhang et al., 2010), the protein coilin is considered to be the CB marker protein. Work conducted by our lab has shown that coilin has RNase activity (Broome et al., 2013), and that this RNase activity produces discrete products when incubated with *in vitro* transcribed scaRNAs (Enwerem et al., 2014). These findings indicate that coilin has function beyond being a building block for CBs. While trying to identify human coilin’s genomic sequence, it was found that there were also two pseudogenes which shared high sequence identity with coilin mRNA: *COILP1* and *COILP2* (Andrade et al., 1993). These pseudogenes contain a number of insertions, deletions and frameshifts. Our lab recently found that *COILP1* has the capacity to encode a protein of 203 residues (Poole et al., 2016). The protein produced by *COILP1* shares sequence homology with coilin’s RNA binding domain (RBD), but has a unique N-terminal sequence. We also found that, unlike coilin, ectopically expressed coilp1 is predominantly localized to nucleoli. It is possible, therefore, that coilp1, along with coilin and other components enriched within the CB may take part in the internal processing of scaRNA 2, 9 and 17.

To further explore the function of the processed fragments derived from scaRNA 2, 9 and 17, and examine if coilp1 plays a part in their formation, we conducted experiments designed to identify *cis* elements within scaRNA 2, 9 and 17 that govern processing and localization. Here we report that the 28 nt sequence at the 5’ end of scaRNA9 plays an important role in stability and protein interactions. We also observed that the level of coilp1 influences the processing of scaRNA 9 and 17. We further show that the stable nucleolus-enriched fragments derived from scaRNA 2, 9 and 17 can base pair with several different snoRNAs. This base pairing may impact snoRNP activity and thereby provide a mechanism by which ribosome heterogeneity is accomplished. We propose that the fragments derived from scaRNA 2, 9 and 17 become novel RNPs that we term regulatory RNPs (regRNPs). Experimental support for the existence of regRNPs is provided, and, if this model for the regulation of ribosome heterogeneity is true, strongly suggest the existence of other regRNPs besides the four we describe here. These findings thus provide a link between splicing and translation by virtue of scaRNA 2, 9 and 17 that modify the splicing machinery when full-length and present in CBs, and regulate rRNA modification when processed into regRNPs that localize to the nucleolus.

## Results

Of the 28 known scaRNAs, only scaRNA 2, 9 and 17 have been shown to be internally processed, which generates stable nucleolus-enriched RNAs of unknown function (Tycowski et al., 2004). Specifically, scaRNA2 gives rise to mgU2-61, scaRNA9 generates mgU2-19 as well as mgU2-30, and scaRNA17 yields mgU4-8 (Fig. 1). Our previous work has shown that the GU repeat region of scaRNA 2 and 9 (shown in Fig. 1) impacts their processing (Enwerem et al., 2015) (Poole et al., 2016). We have also found that coilin associates with scaRNA 2, 9 and 17 (Poole et al., 2016) and reduction of coilin as well as other components of the CB (SMN and WRAP53) differentially affects the processing of scaRNA2 (Enwerem et al., 2015). Since we have found that coilin has nucleic acid processing activity, and this activity is maintained after an extremely stringent purification procedure that greatly reduces the likelihood that the RNA processing activity of coilin is due to co-purifying RNases (Broome and Hebert, 2012), it is possible that coilin directly takes part in some aspect of scaRNA 2, 9 and 17 processing (Broome et al., 2013; Enwerem et al., 2015). Our efforts have also uncovered a novel participant in the formation of RNPs, coilp1, which is encoded by a coilin pseudogene (Poole et al., 2016). We next set out to identify additional *cis* elements and *trans* factors which take part in the processing of scaRNA 2, 9 and 17.

### The scaRNA9 leader sequence is vital for its stability and protein interactions

It was previously reported that scaRNA9 contains, unexpectedly, a 28 nt 5’ leader sequence (Tycowski et al., 2004). Using computational methods, a scaRNA9-like sequence (scaRNA9L) was identified on chromosome X (Yang et al., 2006). It is not known if scaRNA9L, like scaRNA9 (encoded on chromosome 11), is internally processed, but it also appears to have a leader sequence. The presence of the leader sequence strongly suggests that it is playing a functional role, and is being intentionally protected from 5’ exonucleases. Analysis of this leader sequence in scaRNA9 reveals that it contains an AGUG directly before the beginning of the mgU2-19 domain. Interestingly, an AGUG sequence is also found at the beginning of the processed mgU2-61 domain of scaRNA2, immediately before the C box. To examine the role of the scaRNA9 leader sequence, and test the hypothesis that the AGUG might serve as a *cis* element that influences the processing and protein interactions of scaRNA 2 and 9, we generated mutants of these RNAs and conducted RNA pulldown assays. These pulldown assays were conducted using cell lysate and in vitro transcribed biotin labeled RNA. We found that there were no significant changes in the interaction of coilin, SMN, and the known interactor of box C/D RNAs, fibrillarin, with wildtype (WT) scaRNA2 compared to scaRNA2 with a mutated AGUG (Figure 2A, compare lanes 3 to 4). In contrast, deletion of the 28 nt scaRNA9 5’ leader sequence (Δ leader) altered the composition of proteins that associate with this RNA compared to WT: coilin recovery was significantly increased by 2.2-fold yet coilp1 recovery was significantly decreased by 60% (Fig. 2A, compare lane 6 to lane 5, histogram). As positive controls, we show that fibrillarin and SMN recovery does not change, and, as a negative control, we show that there is no β-tubulin recovery by any of the RNAs tested. Moreover, the processing of scaRNA9 without the leader sequence was drastically changed compared to WT (Fig. 2B), as assessed by Northern blotting. Specifically, we observed that, while the levels of the mgU2-30 processed fragment obtained from scaRNA9 Δ leader appear approximately equal to that obtained with WT scaRNA9, the amount of full-length scaRNA9 Δ leader is greatly reduced. We conclude from these findings that the 28 nt leader sequence of scaRNA9 greatly impacts coilin and coilp1 protein binding as well as the stability of full-length scaRNA9. The production of the other fragment derived from scaRNA9, mgU2-19, is also greatly reduced when the 28 nt leader sequenced is deleted (our unpublished observations). These findings clearly show that the 28 nt leader sequence of scaRNA9 is essential for the stability of the full-length scaRNA9 and the mgU2-19 processed fragment, but does not affect the levels of the mgU2-30 fragment. Note that we have previously showed (Enwerem et al., 2015) that the endogenous processed fragments derived from scaRNA 2, 9 and 17 are difficult to detect using Dig probes, necessitating the ectopic expression of these RNAs for detection by Northern.

**Figure 2.**
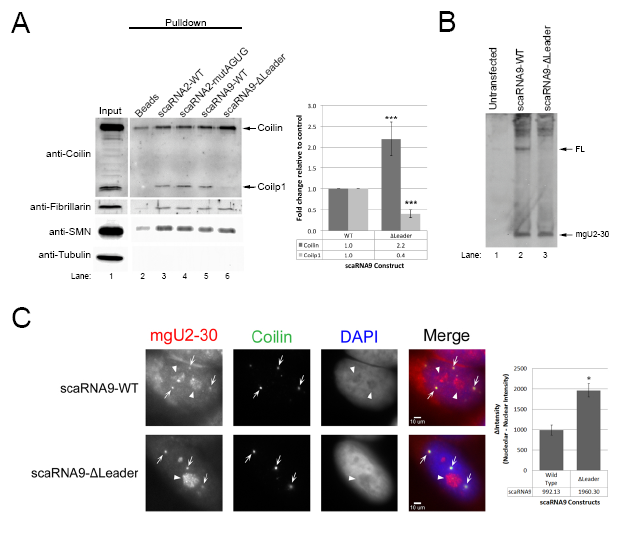
The 28 nt leader sequence of scaRNA9 is important for stability and protein interactions. (A) RNA pulldown of WT and mutant scaRNA2 and scaRNA9. Proteins recovered in the pulldown reactions were subjected to SDS-PAGE, Western transfer, and probing with the indicated antibodies. Background binding of proteins was determined using reactions containing beads alone without RNA bait (lane 2). The input lane (lane 1) accounts for 3% of the lysate used in the pulldown reactions. The location of coilin and coilp1 are indicated, and the relative amount of these proteins recovered by scaRNA9 with the deleted leader sequence (lane 6) compared to WT scaRNA9 (lane 5) is shown in the histogram (n = 5 experimental repeats, ***p < 0.0005, error bars represent SEM). (B) Northern blotting of RNA isolated from untransfected cells (lane 1) or cells transfected with a DNA construct expressing WT scaRNA9 (lane 2) or scaRNA9 with a deleted leader sequence (lane 3). Probe B, shown in Fig. 1, was used to detect full-length scaRNA9 and the mgU2-30 domain. (C) RNA FISH/IF was used to localize WT scaRNA9 and scaRNA9ΔLeader. Probe D, shown in Fig. 1, was used to detect full-length scaRNA9 and the mgU2-30 domain by RNA FISH. Arrows indicate Cajal bodies (detected by coilin localization) and arrowheads demarcate nucleoli (determined by DAPI negative regions). The relative intensity of nucleolar to nucleoplasmic RNA FISH signal was quantified for each RNA (histogram, *p-value < 0.0005, error bars represent SEM, n = 20 cells.

Since the processed fragments derived from scaRNA2, 9 and 17 are enriched in the nucleolus (Tycowski et al., 2004), we next examined the localization of scaRNA9 Δ leader compared to WT scaRNA9. For this work, cells were transfected with DNA encoding WT or Δ leader scaRNA9, and RNA FISH, using a probe indicated by D in Fig. 1, followed by immunofluorescence was conducted (Fig. 2C). Based on the binding location of probe D, full-length scaRNA9 as well as the processed mgU2-30 domain will be detected by this probe. As expected, WT scaRNA9 is enriched in CBs, nucleoli and nucleoplasm (Fig. 2C), consistent with the full-length scaRNA9 being localized in the nucleoplasm/CB and the processed mgU2-30 domain being nucleolar. CBs were detected by anti-coilin staining. The scaRNA9 Δ leader RNA was also detected in CBs and nucleoli, but the intensity of the nucleolar staining relative to that found in the nucleoplasm was significantly increased for the scaRNA9 Δ leader compared to the WT scaRNA9 signal (histogram). These localization data support the Northern results and strongly indicate that the 28 nt 5’ leader sequence of scaRNA9 is important for the stability of full-length scaRNA9. In the absence of this leader sequence, most of the scaRNA9 transcribed from the plasmid DNA is processed into the mgU2-30 domain, which results in greater levels of nucleolar signal compared to that observed with WT scaRNA9.

### Coilp1 positively influences the processing of scaRNA 9 and 17

We have previously shown that *COILP1*, a pseudogene, encodes a 203 amino acid protein which contains sequence homology with coilin’s RNA binding domain, but has an unique 77 amino acid N-terminal sequence (Poole et al., 2016). We further showed that endogenous coilp1 complexes and bacterially purified coilp1 bind *in vitro* transcribed scaRNA2, scaRNA9, and hTR (Poole et al., 2016). Moreover, we found that GFP and myc tagged coilp1 localizes to the nucleolus. Since scaRNA 2, 9, and 17 give rise to stable processed fragments that localize to the nucleolus (Tycowski et al., 2004), we were interested in determining if coilp1 overexpression or knockdown would, like coilin and SMN (Enwerem et al., 2015), affect the biogenesis and/or stability of these processed fragments. For this work, coilp1 was reduced by RNAi, followed by transfection with DNA encoding scaRNA 2, 9 or 17. To examine the effect of coilp1 overexpression on scaRNA 2, 9 and 17 processing, cells were co-transfected with GFP or GFP-coilp1 and DNA encoding scaRNA 2, 9 or 17. Isolated RNA was then subjected to Northern blotting using the probes indicated in Fig. 1. In order to determine the relative amounts of processing, we took the ratio of processed fragment to full length; or, in the case of scaRNA2, ectopic full-length, which runs higher than the endogenous full-length scaRNA2. We found that reduction of coilp1 expression reduced the relative amount of the scaRNA9-derived processed fragment, mgU2-30, by 53% compared to that detected in RNA from control siRNA treated cells (Fig. 3A). In contrast, overexpression of coilp1 resulted in a 1.64-fold increase in the relative amount of mgU2-30 compared to cells expressing GFP alone (Fig. 3B). The amount of coilp1 did not, however, impact the level of scaRNA2-derived mgU2-61 (Fig. 3CD). Upon examination of the processing of scaRNA17 (which generates mgU4-8), we found that, like scaRNA9, reduction of coilp1 decreases the amount of this processed fragment (by 65%, Fig. 3E) while coilp1 overexpression increases relative mgU4-8 levels (by 2.77-fold, Fig. 3F) compared to control treated cells. Together, these data support the hypothesis that coilp1 is a positive regulator of scaRNA 9 and 17 processing.

**Figure 3.**
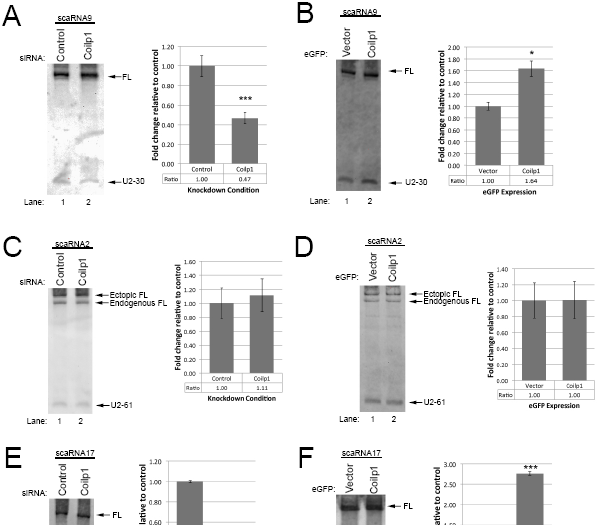
Coilp1 contributes to the processing of scaRNA 9 and 17. For coilp1 overexpression experiments, cells were co-transfected with plasmids expressing scaRNA 2, 9 or 17 with GFP-coilp1 or GFP alone (B, D, F). For coilp1 reduction experiments, cells were treated for 24 hrs with control or coilp1 siRNA, followed by transfection with DNA constructs expressing scaRNA 2, 9 or 17 and harvest/RNA isolation 24 hrs later (A, C, E). The RNA was then subjected to Northern blotting using the probes indicated in Fig. 1. Histograms display the quantification of the processed fragment relative to the full-length scaRNA normalized to the control condition. * p-value < 0.05; *** p-value < 0.0005, error bars represent SEM, n = 3 experimental repeats. A/B, scaRNA9; C/D, scaRNA2; E/F, scaRNA17. Note that ectopic scaRNA9 is expressed from a construct containing an intron. Consequently, full-length ectopic scaRNA9 is the same size as endogenous scaRNA9, which is encoded within an intron. In contrast, full-length ectopic scaRNA2 is longer than endogenous scaRNA2 (Enwerem et al., 2015). Full-length endogenous scaRNA17 is difficult to detect using Dig probes.

### The nucleolus enriched processed fragments of scaRNA 2, 9 and 17: A new class of RNP?

Although it has long been known that the fragments derived from scaRNA 2, 9 and 17 are enriched in the nucleolus, the function of these processed RNAs in this cellular locale is unknown. It was previously thought that these RNAs would guide modification of snRNAs that traffic through the nucleolus (Tycowski et al., 2004). Of the U1, U2, U4, U5 and U6 snRNAs in mammalian cells, however, only the pol III-derived U6 has a clear nucleolar pathway (Kiss, 2004). Since the guide RNAs derived from scaRNA 2, 9 and 17 (mgU2-61, mgU2-19, mgU2-30 and mgU4-8) are thought to act upon U2 and U4, not U6, this leaves open the question as to what these RNAs are doing in the nucleolus. It seems highly unlikely that these RNAs are non-functional by-products given their stability, which strongly indicates the RNAs are part of an RNP. We initially postulated that scaRNA 2, 9 and 17-derived RNAs become snoRNPs, and aid in the modification of rRNA or the nucleolar-trafficked U6 snRNA. Since the processed fragments are box C/D RNAs (and thus guide methylation modifications), we queried several websites, such as snoSCAN (Schattner et al., 2005), that predict target sites for scaRNA 2, 9 and 17 derived RNAs on rRNA. Although these websites returned alignments and methylation target sites on rRNA, these locations did not correspond to any sites with experimentally verified methylation. This lead us to question if these processed RNAs do, in fact, directly modify rRNA. In the process of doing BLAST searches of the scaRNA17-derived mgU4-8 using a snoRNA/scaRNA database (Lestrade et al., 2006), we observed that the 3’ loop region of mgU4-8 can base pair with the snord16 (U16) snoRNA (Fig. 4, bottom right). The snord16 snoRNP guides the methylation of the A484 site of 18S rRNA. We hypothesize that the interaction of mgU4-8 with snord16 might interfere with the association of the snord16 snoRNP with 18S rRNA, thereby inhibiting the methylation A484. In the nucleolus, therefore, mgU4-8 may be part of a new class of RNPs, which we term regulatory RNPs. The key feature of these putative regulatory RNPs is that they are not directly involved in the modification of target sites within the nucleolus, but instead regulate the activity of snoRNPs by interacting with the RNA component of the snoRNP, or the target RNA. Additionally, we further hypothesize that both methylation and pseudouridylation can be regulated by the actions of regulatory RNPs. As shown in Fig. 4, each of the 4 RNAs derived from scaRNA 2, 9 and 17 likely become part of a regulatory RNP that may influence the modification of four sites within 28S rRNA, two sites within 18S rRNA, and two sites within U6 snRNA (Table 1). Regarding U6 snRNA, the 5’ loop region of mgU4-8, intriguingly, contains 8 bases that are found in U6 snRNA, and this loop can base pair with U94 and HBII-166. U94 and HBII-166 guide the methylation of U6 snRNA at C62 and C60, respectively.

**Figure 4.**
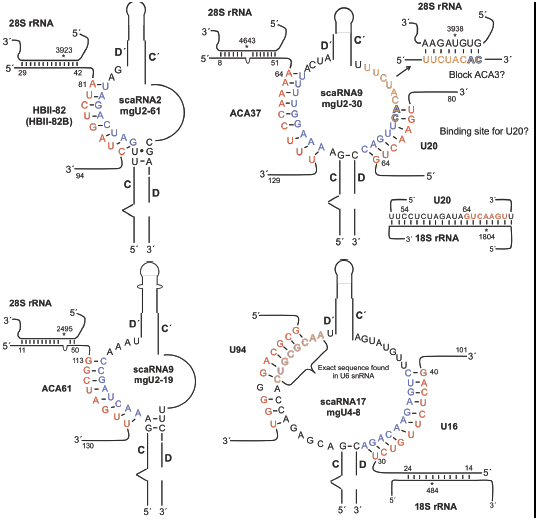
Regulatory RNPs. Base pairing between the nucleolus-enriched fragments of scaRNA 2, 9 and 17 with snoRNA. Also shown is the potential for the 3’ loop of mgU2-30 (top right) to base pair with 28S rRNA, possibly blocking ACA3 binding to this site, as well as serve as a binding site for U20 snoRNA. The 5’ loop of scaRNA17-derived mgU4-8 (bottom right), which contains a sequence found in U6 snRNA, might serve as a binding sink for U94 and HBII-166 (not shown) snoRNAs, thereby altering C60 and C62 methylation of U6 snRNA. We propose that these fragments derived from scaRNA 2, 9 and 17 form regulatory RNPs (regRNPs) that influence the modification of sites within 18S rRNA, 28S rRNA and the nucleolar trafficked U6 snRNA (Table 1).

**Table 1.** Predicted nucleolar targets of scaRNA 2, 9 and 17-derived fragments (regRNPs)

| scaRNA | Processed<br>Fragment | Predicted nucleolar target sites |  |  |
| --- | --- | --- | --- | --- |
|  |  | 28S | 18S | U6 snRNA |
| scaRNA2 | mgU2-61 | G<sub>m</sub>3923 |  |  |
| scaRNA9 | mgU2-19 | Ψ2495 |  |  |
| scaRNA9 | mgU2-30 | Ψ3938, Ψ4643 | U<sub>m</sub>1804 |  |
| scaRNA17 | mgU4-8 |  | A<sub>m</sub>484 | C<sub>m</sub>60, C<sub>m</sub>62 |

### Reduction of scaRNA17 increases the level of A484 methylation within 18S rRNA

To garner experimental support for the existence of regulatory RNPs, cells were treated with control or scaRNA17 siRNAs. In a typical experiment, RNAi reduced scaRNA17 levels by 60% as assessed by qRT-PCR. RNA isolated from these cells was subjected to methylation analysis using a primer extension technique that takes advantage of the fact that reverse transcriptase pauses at sites of methylation when dNTP levels are low (Maden et al., 1995). Primers were designed to interrogate the methylation status of the A484 site of 18S rRNA. The A484 site of 18S rRNA is known to be modified, presumably due to association with the snord16 box C/D snoRNP. Since the mgU4-8 fragment derived from scaRNA17 is enriched in the nucleolus and has extensive base pairing with snord16 snoRNA (Fig. 4), and this interaction may inhibit snord16 interaction with 18S rRNA, we predicted that scaRNA17 reduction (which would lead to the reduction of mgU4-8) would increase the methylation level of A484 within 18S rRNA. This is exactly what we have observed (Fig. 5). RNA obtained from cells in which scaRNA17 was reduced had consistently more (1.8-fold) A484 methylation compared to that observed in RNA from control siRNA treated cells (n = 3, *P*<0.05). In contrast, 2’-O-methylation levels were essentially the same at the other two known 2’-O-methylation sites (Am436 and Am468), which are not guided by snord16 or targeted by mgU4-8 (derived from scaRNA17). We also observed an increase in the pause site (denoting increased methylation) of the A484 site in scaRNA17-reduced RNA using a primer labeled with digoxigenin (Fig. 5B). These results thus argue that the processed fragment derived from scaRNA17 can form a regulatory RNP that downregulates the methylation of the A484 site in 18S rRNA.

**Figure 5.**
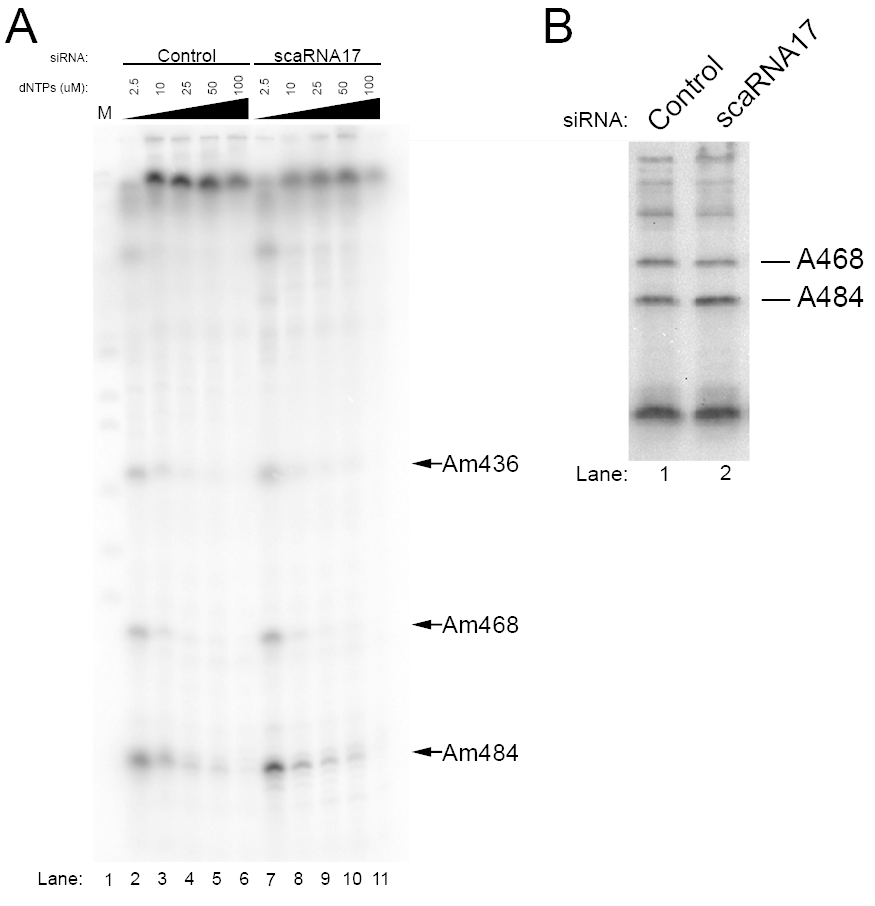
Methylation of 18S A484 is increased upon reduction of scaRNA17.

(A) RNA from control or scaRNA17 siRNA treated cells was subjected to primer extension using 5 different dNTP amounts (indicated) during the reverse transcription step with a radioactive primer complementary to nucleotides 524-544 of 18S rRNA. Samples were run on a denaturing acrylamide gel followed by detection of the radioactive signals. (B) Primer extension was conducted using only low (2.5 uM) dNTP and the primer is 5’ Dig labeled. Quantification demonstrates that the A484 pause signal is 1.8-fold more abundant in the scaRNA17 knockdown condition compared to control knockdown (n = 3 experimental repeats, p < 0.05).

### Disruption of snord16/18S rRNA interaction by a fragment of scaRNA17

Previous work has shown that methylation of rRNA by some box C/D snoRNPs is facilitated by “extra base pairings” between the snoRNA and target rRNA (van Nues et al., 2011). Most of these extra base pairings are the result of loops within the snoRNA, allowing for additional snoRNA:rRNA interactions. The interaction between snord16 snoRNA with 18S rRNA is an example of an association that contains extra base pairings (Fig. 6A). An additional 10 base pairings between snord16 snoRNA and 18S rRNA is made possible by a loop of snord16, between nucleotide 24 and 48. In so doing, it is expected that the methylation of A484 of 18S rRNA is increased as a consequence of these extra base pairings compared to the level of methylation if only nucleotides (nt) 14-24 of snord16 base paired with 18S rRNA. Very interestingly, the scaRNA17-derived nucleolar fragment mgU4-8 interacts with snord16 via nucleotides present in the snord16 loop region (Fig. 6B). It is possible, therefore, that the association of mgU4-8 with the looped region of snord16 disrupts the interaction between snord16 with 18S rRNA, resulting in a decrease in the level of 18S rRNA A484 methylation. Given this regulatory activity, we propose that the mgU4-8 domain generated by processing of scaRNA17 be renamed regulatory RNP17 (regRNP17).

**Figure 6.**
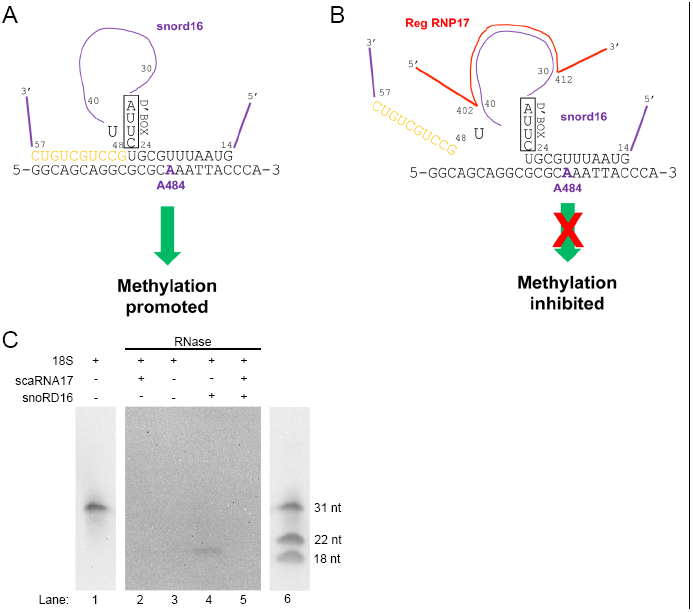
Disruption of the snord16/18S rRNA interaction by a fragment of scaRNA17. (A) Extra base paring (shown in orange) (van Nues et al., 2011) between snord16 and 18S rRNA predicted to facilitate the methylation of A484 (shown in purple) is shown. (B) The looped region of snord16 (from nt 30-40) base pairs with scaRNA17 (regRNP17) from nt 412-402 (Fig. 4). The binding of the nucleolus-enriched fragment derived from scaRNA17, mgU4-8 (regRNP17), may disrupt the interaction of snord16 with 18S rRNA, resulting in decreased A484 methylation. (C) RNase protection assays using fragments of snord16, 18S rRNA (31 nt, 3’ Dig labeled) and scaRNA17. Reactions containing the indicated RNA were denatured, followed by annealing and incubation with RNase A/T1. The reactions were then resolved on a 15% TBE-urea polyacrylamide gel, followed by Northern transfer and detection with anti-Dig antibodies. Because of the intensity of the signal, only 20% of the reaction with 18S rRNA fragment lacking RNase A/T1 was run on the gel (lanes 1 and 6), and the exposure of these lanes is shorter than that for lanes 2-5. Additionally, the reaction sample in lane 6 was supplemented with Dig-labeled DNA oligonucleotides to serve as size markers (indicated by bands at 22 and 18 nt). A protected fragment is observed in lane 4 but not in lane 5.

To begin an analysis into the mechanism by which regRNP17 imparts a regulatory effect upon the modification of rRNA, we conducted in vitro RNase protection assays using fragments of 18S rRNA, U16 snoRNA and scaRNA17. The 18S rRNA fragment is 31 nt in length, contains the A484 site and is 3’ end labeled with digoxigenin (Dig). The snord16 fragment is 45 nt in length and encompasses nucleotides 14-57 (Fig. 6). The regRNP17 fragment is 33 nt in length and contains the bases that interact with snord16. The basis for RNase protection is that interacting RNAs will not be subjected to degradation by RNase A/T1, which preferentially cleaves single stranded RNA. Consequently, we expect that the interaction of the snord16 fragment with the 18S fragment will result in a protected fragment approximately 22 nt in length. The addition of the regRNP17 fragment to the 18S/snord16 mixture, however, is predicted to disrupt the interaction between snord16 and 18S, decreasing the amount of Dig-labeled 18S fragment that is protected from RNase A/T1 degradation. This is what we have observed (Fig. 6C). In a reaction containing just the Dig-labeled 18S rRNA fragment incubated without RNase A/T1 (lanes 1 and 6), a 31 nt band is detected. This band is digested upon incubation with RNase A/T1 (lane 3). The addition of snord16 to the reaction with the 18S rRNA results in a protected fragment (lane 4), indicating that the interaction between snord16 and 18S precludes RNase A/T1 from fully digesting the 18S RNA. In a reaction containing all three fragments (18S, snord16 and regRNP17), however, the amount of 18S protected fragment is decreased (lane 5), suggesting that the regRNP17 disrupts the interaction between snord16 and 18S rRNA. A reaction containing regRNP17 and 18S does not result in a protected fragment (lane 2), demonstrating that snord16 is required to generate the protected fragment. These findings strongly suggest that regRNP17 influences A484 methylation levels by altering the interaction between snord16 and 18S rRNA.

## Discussion

### A novel RNP: the regulatory RNP

In 2004 it was found that scaRNA 2, 9 and 17 are internally processed, resulting in the generation of stable nucleolus-enriched fragments (Tycowski et al., 2004). Base pairing between the full-length scaRNA 2, 9 and 17 with U2, U4 and U12 snRNAs strongly suggest that scaRNA 2, 9 and 17 guide the 2’-*O*-methylation of specific sites within these snRNAs. Like other scaRNAs, therefore, full-length scaRNA 2, 9 and 17 contribute towards the biogenesis of snRNPs, which are crucial parts of the spliceosome necessary for pre-mRNA splicing. The function of the stable, nucleolus-enriched fragments derived from these three scaRNAs, however, remains murky. Specifically, the function of mgU2-61 (derived from scaRNA2), mgU2-19 and mgU2-30 (both derived from scaRNA9) and mgU4-8 (derived from scaRNA17) (Fig. 1) is not evident considering that their putative targets (U2 and U4 snRNA) do not have a clear nucleolar pathway (Kiss, 2004). By conducting BLAST searches of the unpaired loops of these processed fragments against a snoRNA database (Lestrade et al., 2006), we found that each of the four fragments derived from scaRNA 2, 9 and 17 can base pair with snoRNA (Fig. 4). This led us to speculate that the processed fragments do not form RNPs which directly methylate targets, but instead may regulate the activity of snoRNPs responsible for 2’-*O*-methylation and pseudouridylation (Fig. 4, Table 1). Experimental evidence in support of this hypothesis is shown in Figs. 5 and 6, and argues for the possibility that the nucleolar-enriched processed fragments of scaRNA 2, 9 and 17 form novel RNP complexes that we term regulatory RNPs (regRNPs). We propose that regRNPs affect the modification of rRNA and U6 snRNA (which traffics though the nucleolus) by interactions with the snoRNA component of snoRNPs. It is highly unlikely that there are only four regRNPs present (one each from scaRNA 2 and 17 and two from scaRNA9). Rather, we believe that some regRNPs are incorrectly classified as snoRNPs or orphan snoRNPs. It is conceivable, therefore, that there are numerous regRNPs which work together to ensure that the level of rRNA modifications are appropriate for a given situation.

### RegRNPs and ribosome heterogeneity

An exciting emerging concept is that of ribosome heterogeneity, leading to specialized ribosomes. This concept is based on the realization that ribosomes are not all the same, all the time, but can vary in response to different physiological or pathological situations (Lafontaine, 2015). One major contributor to ribosome heterogeneity is rRNA modification. Since each ribosome in human contains around 100 each of pseudouridines and ribose methylations, there is a vast potential for ribosome specialization in regulating the level of these modifications in rRNA (Lafontaine, 2015). Very significantly, three methylation sites within rRNA that we predict are subject to regRNP 2, 9 and 17 control (G3923 in 28S, U1804 and A484 in 18S) have been shown to be differentially modified in endogenous ribosomes (Krogh et al., 2016) (Incarnato et al., 2017). In fact, all three of these sites show upwards of 25% variability (Krogh et al., 2016) (Incarnato et al., 2017). Interestingly, comparison of the U1804 and G3923 methylation levels in HeLa and HCT116 cells shows significant differences between these lines (Krogh et al., 2016). These differences were not correlated with altered levels of snoRNPs, indicating that another factor besides snoRNP availability is responsible for the differential modification of rRNA (Krogh et al., 2016). Our data presented in Figs. 5 and 6 showing that regRNP17 impacts the level of A484 methylation in 18S rRNA supports the hypothesis that ribosomes may be optimized for a given cell type or physiological/pathological situation by controlling the level of rRNA modifications using regRNPs.

### 18S rRNA/snord16/regRNP17 interactions in context with other snoRNAs

Given that rRNA has extensive secondary structure as well as numerous modifications, it is not surprising that the accessibility and regulation of snoRNP activity must be subject to some type of control. An example of where one would expect some type of ordered snoRNP activity is shown in Fig. 7A, which displays the snoRNAs that interact with 18S rRNA in the region of A484. In particular, snord11, snord56 and snord70 have overlapping binding sites on the 18S rRNA. Moreover, a 9 nt region of 18S rRNA starting at T514 (underlined in Fig. 7A) is found exactly in snord16. In fact, snord16 nt 76-86 can complementary base pair with snord56 at nt 29-19. In so doing, interactions between snord16 and snord56 may contribute towards the regulation of C517 methylation. Since regRNP17 interacts with the looped region of snord16, and this association appears to disrupt the interaction of snord16 with 18S rRNA (Fig. 6), it is further possible that regRNP17 may also be involved in the regulation of snord56-mediated C517 methylation. Snord16 has many RNA interactions that may regulate its functions, and, conversely, allow snord16 to regulate the activities of the RNAs it interacts with (Fig. 7B). In fact, 65% of snord16 can base pair with other RNAs (66 nt out of 101 nt). A key regulator of these snord16 interactions may be regRNP17, which, by binding the looped region of snord16, could greatly influence snord16 associations with 18S rRNA and other snoRNAs.

**Figure 7.**
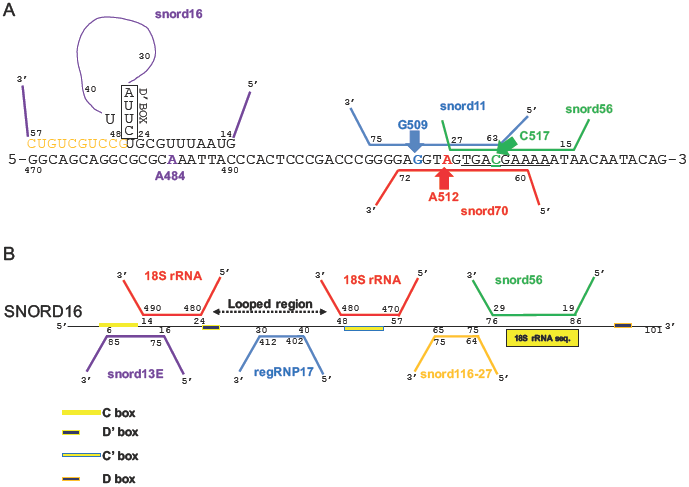
The A484 region of 18S rRNA, and snord16 associated RNAs. (A) A schematic of 18S rRNA is shown, along with interacting snoRNAs. Note that snord16 can form extra base pairing with 18S rRNA (orange nt) by the formation of a loop within snord16 (van Nues et al., 2011). Modification of G509, A512 and C517 involves guide snoRNAs that having overlapping binding site. The underlined sequence in 18S rRNA including C517 is found exactly in snord16, which means that snord56 can associate with snord16. (B) Schematic of snord16, which is 101 nt long. The looped region of snord16, which interacts with regRNP17, is indicated. The sequence in snord16 that is exactly identical to that found in 18S rRNA is indicated by a yellow rectangle. The locations of the snord16 C/D and C’/D’ boxes are shown, as well as associations with other snoRNAs.

### Identification of a *cis* element in scaRNA9

A limitation to the understanding of why scaRNA 2, 9 and 17 are internally cleaved has been the identification of *cis* elements within these RNAs that govern their processing. We have previously observed that the GU rich repeat regions in scaRNA 2 and 9 (Fig. 1) impacts their processing (Enwerem et al., 2015; Poole et al., 2016), and here identify another *cis* element: the 28 nt leader sequence of scaRNA9 (Fig. 2). We have shown that the 28 nt leader sequence of scaRNA9 is critical for its stability and influences its protein interactions. Since scaRNA9 is processed at its 5’ and 3’ end by exonucleases, any nucleic acid sequence not protected by a protein complex should be processed. This indicates that there are protein components which bind to the scaRNA9 leader sequence that block it from being processed. We show here that coilp1 could be one of these interactors, as coilp1’s interaction with scaRNA9 Δ leader is drastically reduced compared to WT scaRNA9 (Fig. 2A). Furthermore, the increased recovery of coilin (2.2-fold) using the scaRNA9 Δ leader bait in the RNA pulldown compared to the amount of coilin obtained with WT scaRNA9 may indicate that coilp1 negatively regulates the amount of coilin in the binding complex. Not surprisingly, only full-length scaRNA9 and the 5’-most processed fragment (mgU2-19) levels are deleteriously impacted by deletion of the leader sequence (Fig. 2B). An interesting future direction involves deciphering the mechanism which governs the amount of scaRNA 2, 9 and 17 that is processed into regRNPs.

### Coilp1 contributes to the availability of regRNPs

The data presented here further characterizes the recently discovered pseudogene encoded protein coilp1, and implicates it in the biogenesis of the newly described regRNPs. At this point, the exact function of coilp1 in this process is not clear. ScaRNA 9 and 17 showed an increase in processing when coilp1 was overexpressed, and a decrease in processing when coilp1 expression is reduced (Fig. 3). However, we did not observe any change in processing for scaRNA2. ScaRNAs 2 and 17 are both independently transcribed by RNA polymerase II (Gerard et al., 2010; Tycowski et al., 2004). Possibly due to non-canonical box C and D sequences, mgU2-25 (scaRNA2) and mgU12-22 (scaRNA17) are inherently unstable, and do not accumulate within the cell once the scaRNAs are internally processed. ScaRNA9 is contained within a host gene, and, as such, must be spliced and processed at its 5’ and 3’ ends. ScaRNA9’s 5’ domain, mgU2-19, is capable of accumulating, albeit a lower levels than its 3’ domain, mgU2-30. Indeed, work previously done by our lab has shown that scaRNA2 is less efficiently processed by coilin than scaRNA9 (Enwerem et al., 2015). Given that coilp1 shares sequence homology with coilin, coilp1’s differential contribution to the availability of regRNPs may indicate the existence of further specialization within each scaRNA classification. In summary, based upon the data presented here and our previous work (Enwerem et al., 2015) (Poole et al., 2016), we believe that coilin, WRAP53, SMN and coilp1 all participate at regulating the flux of scaRNA 2, 9 and 17 distribution and processing. We further predict that coilin negatively regulates the activity of coilp1 but SMN and WRAP53 promote this activity (Fig. 8). Another possibility is that coilp1 may chaperone regRNPs to the nucleolus. Given that Nopp140 is a snoRNP chaperone, interacts with coilin and localizes to the CB and nucleolus (Isaac et al., 1998), it is also well positioned to serve as an important factor in the formation of regulatory RNPs. Indeed, deletion of Nopp140 in fly reduces rRNA methylation (He et al., 2015), supporting a role for this protein in the formation of box C/D RNPs. Future work will thus explore the role of Nopp140 on the biogenesis of regRNPs.

In conclusion, our work has provided a possible function for the processed fragments derived from scaRNA 2, 9 and 17 as regRNPs. Although not an exact parallel to what we are proposing, the Stamm group has shown that box C/D snoRNAs can have dual functions in rRNA modification and alternative pre-mRNA splicing (Falaleeva et al., 2016). Thus there is some degree of precedent for a guide RNA having additional functions. Our concept of a regulatory RNP, however, is novel. These regRNPs are predicted to contribute to the heterogeneity of RNA, leading to specialized ribosomes. These findings further add to the complexity of ribosome biogenesis. When ribosome biogenesis is negatively affected, it results in the ribosomopathy disease state, which is characterized by any dysfunction in any of the hundreds of components which facilitate ribosome formation (Wang et al., 2015; Zhou et al., 2015). Interestingly, recent work in human cells has shown that p53 down regulates fibrillarin levels, and in cancer cells lacking functional p53 the level of rRNA methylation is increased (Marcel et al., 2013). This increase in rRNA methylation results in ribosomes with a lower fidelity (i.e. stop codons are bypassed) and a greater likelihood to initiate translation through internal ribosome entry sequences (IRESs) (Belin et al., 2009; Marcel et al., 2015; Marcel et al., 2013). As a consequence of these changes in rRNA methylation, the translation of messages with IRESs is increased; such as those whose products promote tumor development (IGF-1R, c-myc, VEGF-A and FGF1) (Marcel et al., 2015). Clearly, therefore, the regulation of modifications within rRNA is of great importance. With our identification of regRNPs, we demonstrate that one non-coding RNA can, based on its localization and processing, influence both the splicing and translation machinery. Our current efforts seek to experimentally verify additional regRNPs.

**Figure 8.**
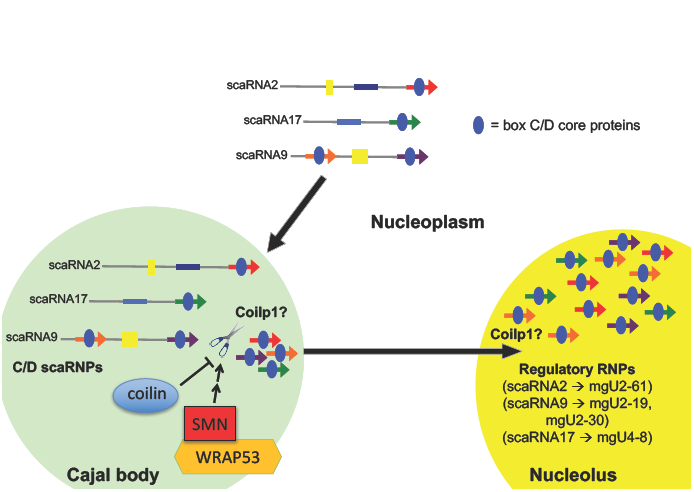
Model of box C/D scaRNP 2, 9 and 17 biogenesis. Box C/D core proteins (blue) bind scaRNAs after transcription from independent genes (scaRNA2 and scaRNA17) or host gene (scaRNA9). GU dinucleotide repeats (yellow box) in scaRNA2 and 9 are indicated. Nucleolus-enriched RNAs derived from scaRNA2 (mgU2-61), scaRNA9 (mgU2-19 and mgU2-30) and scaRNA17 (mgU4-8), which we propose form regulatory RNPs, are shown. Coilin, SMN and WRAP53 may regulate the activity of Coilp1. It is possible that these interactions take place in the CB (or the nucleoplasm in cells lacking CBs). It is also possible that coilp1 participates in the transport of regRNPs to the nucleolus. Note that the color coding here is the same as that in Fig. 1.

## Materials and Methods

### Cell lines, cell culture, plasmids, and transfections

HeLa cells were cultured as previously described (Enwerem et al., 2014). EGFP-C1 (Clontech, Mountain View, CA, USA) was used for empty vector experiments. GFP-coilp1 (Poole et al., 2016), scaRNA2 pcDNA 3.1+ (Enwerem et al., 2014), and scaRNA9 pcDNA3.1+ (Enwerem et al., 2015) have been previously described. ScaRNA17 was amplified from genomic DNA isolated from HeLa-ATCC using standard molecular biology techniques, and cloned into pcDNA3.1+ using *Bam*H1 and *Eco*RI restriction sites. Primers used to amplify scaRNA17: Forward (5’-AGAGGCTTGGGCCGCCGAGCT-3’) Reverse (5’-TCTGAGAACAGACTGAGGCCG-3’). ScaRNA2-mut AGUG and scaRNA9ΔLeader were generated from the aforementioned pCDNA3.1+ wild type clones using site directed mutagenesis. The following primers were used: scaRNA2-mutAGTG: Forward (5’-TGCGGGGCCCGGCGCTCAGATCACATGAATTGATCAGATAGACG-3’) Reverse (5’-CGTCTATCTGATCAATTCATGTGATCTGAGCGCCGGGCCCCGCA-3’) scaRNA9ΔLeader: Forward (5’-ATCGTCGCAGGATCCGATCAATGATGAAACTAGCC-3’) Reverse (5’-GGATCCTGCGACGATGCACTGACTTTAATGTTATAAC-3’). To generate *in vitro* transcribed RNAs, scaRNA2, scaRNA2-AGTG, scaRNA9, and scaRNA9ΔLeader were inserted into pBluescript KS vectors using the same restriction sites from the pCDNA3.1+ clones, and were transcribed using MEGAscript T7 (Thermo Fisher, Waltham, MA, USA) (scaRNA9ΔLeader) or MEGAscript T3 kit (Thermo Fisher, Waltham, MA, USA) (scaRNA2, scaRNA2-AGTG, and scaRNA9) with 40% Biotin-UTP (Roche, Mannheim, Germany). DNA transfections were done using FuGene (Promega, Madison, WI, USA) following the manufacturer’s suggested protocol for 24-hour expression. SiRNA transfections were conducted using either Lipofectamine 2000 or RNAi Max (Invitrogen, Carlsbad, CA, USA) following the manufacturer’s suggested protocol for 48-hour knockdown. Control, coilin, WRAP53, SMN and coilp1 siRNAs were previously described (Poole et al., 2016). scaRNA17 expression was reduced using the following dsiRNA: Sense (5’-CCGCAGUAUUUUCCUUAUAUGAUCA-3’) anti-sense (5’-UGAUCAUAUAAGGAAAAUACUGCGGGC-3’). This dsiRNA resulted in a ˜60% knockdown (n=3, Ρ<.05) as determined by qRT-PCR using a previously described protocol (Poole and Hebert, 2016). Primers used to amplify scaRNA17 for qRT-PCR analysis: Forward (5’-GCTGGACCCGGACCGGTTTTGGG-3’) Reverse (5’-AAGGAAAATACTGCGGGCTCATCC-3’). For combined expression and knockdown experiments, cells were first transfected with siRNA for 24 hours then transfected again with DNA for an additional 24 hours for 48-hour knockdown and 24-hour expression.

### RNA isolation and Northern blot

RNA was isolated using TRI-Reagent (Molecular Research Center Inc., Cincinnati, OH, USA) following the manufacturer’s suggested protocol. RNAs were loaded in Gel Loading Buffer II (Roche, Mannheim, Germany), incubated at 95°C for 5 minutes, and run on a 6% TBE urea gel (6% 19:1 acrylamide:bis-acrylamide, 1× TBE, and 7M urea). Gels were rinsed in 1× TBE (Sigma, St. Louis, MO, USA) for 10 minutes prior to being transferred to DNA transfer stacks (Invitrogen, Carlsbad, CA, USA) using an iBlot (Invitrogen, Carlsbad, CA, USA) on setting P8 for 8 minutes. RNA was crosslinked to the membrane using a UV-crosslinker at 120,000 μJ/cm^2^. Membranes were pre-hybridized using UltraSensitive UltraHyb (Thermo Fisher, Waltham, MA, USA) before overnight incubation in a hybridization oven at 37°C with DIG labeled probes (DIG Tailing Kit, Roche, Mannheim, Germany). The following DNA oligonucleotides (Integrated DNA Technologies, Coralville, Iowa, USA) were used for detection: 5’-AGTGGCCGGGGACAAGCCCGGCCTCGTCTATCTGATCAATTCATCACTTCT-3’ (scaRNA2); 5’ TAGAAACCATCATAGTTACAAAGATCAGTAGTAAAACCTTTTCATCATTGCCC-3’ (scaRNA9); 5’-AACTCAGATTGCGCAGTGGTCTCGTCATCA-3’ (scaRNA17). Membranes were then prepared for processing using the DIG Wash and Block kit (Invitrogen, Carlsbad, CA, USA) following the manufacturer’s suggested protocol. Detection was carried out using CSPD (Roche, Mannheim, Germany) following the manufacturer’s suggested protocol.

### RNA Pulldown, Western blot, and antibodies

RNA pulldowns were conducted as previously described (Poole et al., 2016). The following antibodies were used: anti-coilin (Santa Cruz, Dallas, Texas, USA), anti-fibrillarin (Santa Cruz, Dallas, Texas, USA), anti-SMN (Abcam, Cambridge, UK), and anti-β-tubulin (Santa Cruz, Dallas, Texas, USA). Membranes were visualized and quantified using a ChemiDoc (BioRad, Hercules, CA, USA) with QuantityOne software. Data was imported to Microsoft Excel, and statistical significance determined using the Student’s t-test.

### Primer extension to detect 2’-*O*-methylation

Reverse transcriptase stops one base downstream of methylated residues at low dNTP concentrations and primer extension to detect 2’-*O*-methylation was carried out as described previously (Huang et al., 2016). The primer used to amplify 18S rRNA and designed to interrogate A484 methylation is: 5’-ATTGTTATTTTTCGTCACTAC-3’. This primer was radioactively or digoxigenin labeled at the 5’ end. Reactions were run on an 8% sequencing gel (for radioactive samples) or a 15% TBE-Urea Novex pre-cast gel (Invitrogen, Carlsbad, CA) (for dig labeled samples). Radioactivity was then detected overnight using a phosphorimager, whereas the gels for the dig labeled products were Northern blotting (as described above) followed by detection with anti-dig antibodies (no overnight hybridization step).

### RNase Protection Assay

RNA protection assays were performed using fragments of snord16 (5’-GUAAUUUGCGUCUUACUCUGUUCUCAGCGACAGUUGCCUGCUGUC), scaRNA17 (5’-GAUGGAGUAUGUUCUGAGAACAGACUGAGGCCG-3’), and 3’ DIG-labeled 18S rRNA (5’-AUCCAAGGAAGGCAGCAGGCGCGCAAAUUAC/3Dig_N/-3’) obtained from Integrated DNA Technologies (Coralville, Iowa, USA). Reactions were performed using the RPA III kit (Thermo Fisher, Waltham, MA, USA). 1 μL fragment (10 μM) and up to 9 μL H2O were denatured at 95C for 2 minutes then annealed by cooling to room temperature for 5 minutes. Reactions were then placed on ice, and either 1 μL digestion buffer (RPA III kit) or 1 μL of a 1:250 dilution of RNase A/RNase T1 Mix (RPA III kit) were added. Digestion was then carried out at room temperature for 30 minutes. Protected fragments were then analyzed by Northern blotting as described above for primer extension. 20% of a reaction lacking RNase was used for input. The following oligonucleotides were added to a reaction lacking RNase to serve as size markers: 22 nt (5’-/5DigN/ATTGTTATTTTTCGTCACTACC-3’) and 18 nt (5’-/5DigN/CCTACGGAAACCTTGTTA-3’).

### IF and RNA FISH

Cells were fixed in 4% PFA for 10 minutes and permeabilized in 0.5% Triton for 5 minutes. Slides were then rinsed 3 times in 1× PBS. Slides were blocked using 10% normal goat serum (NGS) for 30 minutes at 37°C. Slides were then probed with 10% NGS containing 1:200 coilin antibody for 30 minutes at 37°C. Slides were then washed 3 times for 5 minutes, and incubated with 1:600 AlexaFlour-488 Goat anti-rabbit secondary antibody in 10% NGS for 30 minutes at 37°C. Slides were then washed 5 times for 5 minutes in 1× PBS. Following immunostaining, cells were post-fixed in 4% PFA for 10 minutes. Slides were then washed twice in 2× SSC for 5 minutes followed by two washes in 70% ethanol. Cells were then dehydrated by a series of ethanol washes (80%, 95%, and 100% ethanol) for 3 minutes each, and slides were allowed to air dry for 5 minutes. 100 μl of probe solution (10% dextran sulfate, 2 mM VRC, 0.02% BSA, 40 μg *E. coli* tRNA, 2× SSC, 50% formamide, and 30 ng probe) was added to the slides and incubated overnight at 37°C. Probe used for detection of scaRNA9 (Integrated DNA Technologies, Coralville, Iowa, USA): 5’-/5Alex594N/TCATAGTTACAAAGATCAGTAGTAAAACCTTTTCATCATTG-3’. Slides were then washed 3 times for 5 minutes each in 2× SSC, and DAPI stained for 5 minutes. Slides were destained in 2× SSC for 5 minutes prior to mounting.

### BLAST searches

Using a publicly available database (Lestrade and Weber, 2006), BLAST were carried out by using sequences from scaRNAs 2, 9, and 17 that were previously reported to be contained within loop structures (Tycowski et al., 2004). Antisense hits were then compared to reported structures of predicted targets, where applicable, to validate the accessibility of the complementary region.

## Acknowledgments

This work was supported by the Intramural Research Support Program of the University of Mississippi Medical Center (to MDH) and NIH grant GM104077 (to YTY).

---

**Supplemental Figure 1.**
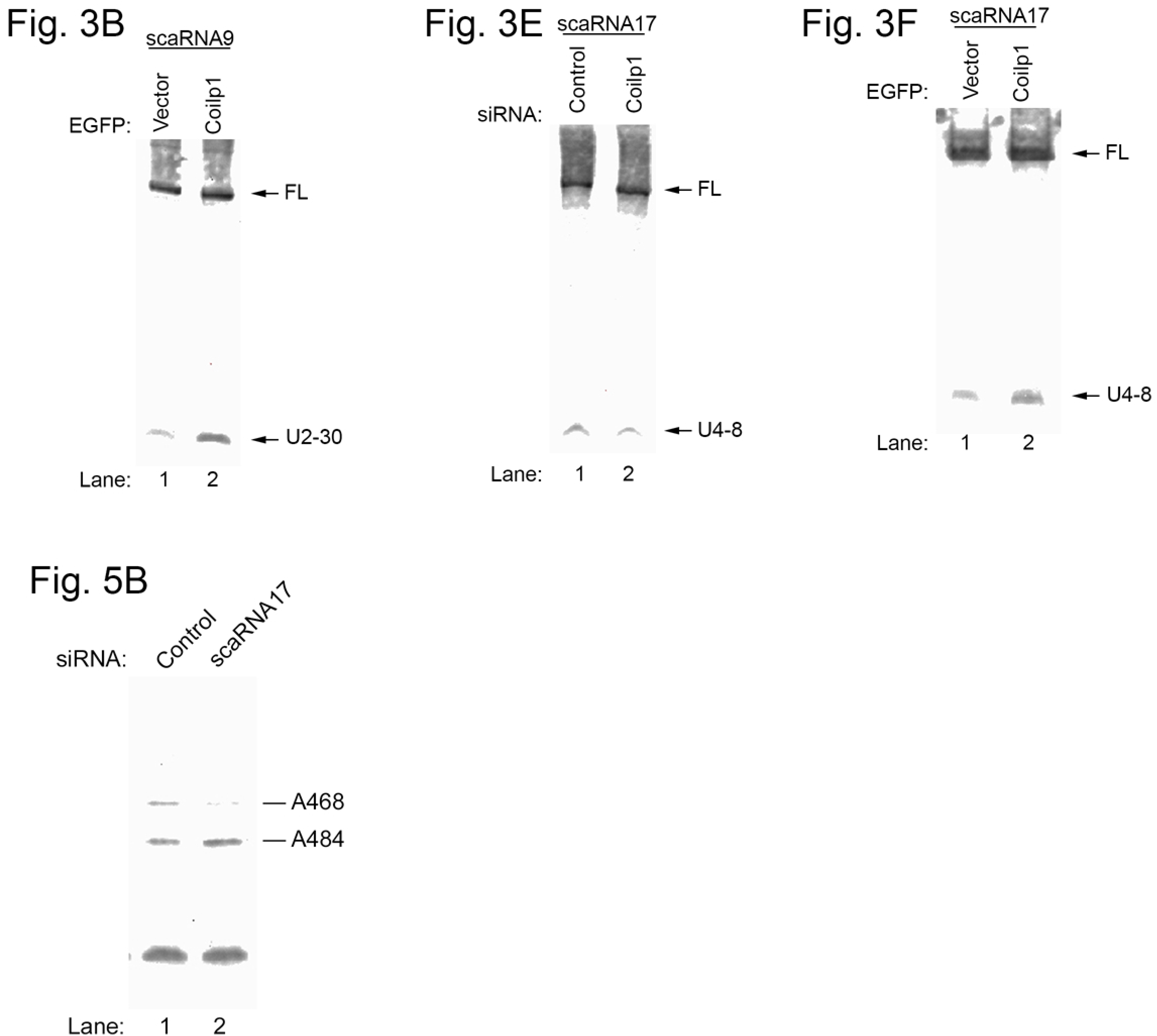
Transformed images. In order to more easily observe the differences between the control and experimental conditions, the images for Fig. 3B, Fig. 3E, Fig. 3F and Fig. 5B have been transformed using the QuantityOne software. Only High and Low transformation settings were adjusted; gamma levels were not changed. This transformation was applied evenly across the entire image for each individual panel.

## References

Andrade, L. E., Tan, E. M. and Chan, E. K. (1993). Immunocytochemical analysis of the coiled body in the cell cycle and during cell proliferation. Proc Natl Acad Sci U S A 90, 1947–51.

Balakin, A. G., Smith, L. and Fournier, M. J. (1996). The RNA world of the nucleolus: two major families of small RNAs defined by different box elements with related functions. Cell 86, 823–34.

Baserga, S. J., Yang, X. D. and Steitz, J. A. (1991). Three pseudogenes for human U13 snRNA belong to class III. Gene 107, 347–8.

Belin, S., Beghin, A., Solano-Gonzalez, E., Bezin, L., Brunet-Manquat, S., Textoris, J., Prats, A. C., Mertani, H. C., Dumontet, C. and Diaz, J. J. (2009). Dysregulation of ribosome biogenesis and translational capacity is associated with tumor progression of human breast cancer cells. PLoS One 4, e7147.

Bousquet-Antonelli, C., Henry, Y., G’Elugne J, P., Caizergues-Ferrer, M. and Kiss, T. (1997). A small nucleolar RNP protein is required for pseudouridylation of eukaryotic ribosomal RNAs. EMBO J 16, 4770–6.

Broome, H. J., Carrero, Z. I., Douglas, H. E. and Hebert, M. D. (2013). Phosphorylation regulates coilin activity and RNA association. Biol Open 2, 407–15.

Broome, H. J. and Hebert, M. D. (2012). In vitro RNase and nucleic acid binding activities implicate coilin in U snRNA processing. PLoS One 7, e36300.

Darzacq, X., Jady, B. E., Verheggen, C., Kiss, A. M., Bertrand, E. and Kiss, T. (2002). Cajal body-specific small nuclear RNAs: a novel class of 2’-O-methylation and pseudouridylation guide RNAs. Embo J 21, 2746–56.

Enwerem, II, Velma, V., Broome, H. J., Kuna, M., Begum, R. A. and Hebert, M. D. (2014). Coilin association with Box C/D scaRNA suggests a direct role for the Cajal body marker protein in scaRNP biogenesis. Biol Open 3, 240–9.

Enwerem, II, Wu, G., Yu, Y. T. and Hebert, M. D. (2015). Cajal body proteins differentially affect the processing of box C/D scaRNPs. PLoS One 10, e0122348.

Falaleeva, M., Pages, A., Matuszek, Z., Hidmi, S., Agranat-Tamir, L., Korotkov, K., Nevo, Y., Eyras, E., Sperling, R. and Stamm, S. (2016). Dual function of C/D box small nucleolar RNAs in rRNA modification and alternative pre-mRNA splicing. Proc Natl Acad Sci U S A 113, E1625–34.

Fatica, A., Galardi, S., Altieri, F. and Bozzoni, I. (2000). Fibrillarin binds directly and specifically to U16 box C/D snoRNA. RNA 6, 88–95.

Ganot, P., Caizergues-Ferrer, M. and Kiss, T. (1997). The family of box ACA small nucleolar RNAs is defined by an evolutionarily conserved secondary structure and ubiquitous sequence elements essential for RNA accumulation. Genes Dev 11, 941–56.

Gautier, T., Berges, T., Tollervey, D. and Hurt, E. (1997). Nucleolar KKE/D repeat proteins Nop56p and Nop58p interact with Nop1p and are required for ribosome biogenesis. Mol Cell Biol 17, 7088–98.

Gerard, M. A., Myslinski, E., Chylak, N., Baudrey, S., Krol, A. and Carbon, P. (2010). The scaRNA2 is produced by an independent transcription unit and its processing is directed by the encoding region. Nucleic Acids Res 38, 370–81.

He, F., James, A., Raje, H., Ghaffari, H. and DiMario, P. (2015). Deletion of Drosophila Nopp140 induces subcellular ribosomopathies. Chromosoma 124, 191–208.

Hebert, M. D. and Poole, A. R. (2016). Towards an understanding of regulating Cajal body activity by protein modification. RNA Biol, 0.

Huang, C., Wu, G. and Yu, Y. T. (2016). Purification and Functional Reconstitution of Box H/ACA Ribonucleoprotein Particles. Methods Mol Biol 1421, 97–109.

Incarnato, D., Anselmi, F., Morandi, E., Neri, F., Maldotti, M., Rapelli, S., Parlato, C., Basile, G. and Oliviero, S. (2017). High-throughput single-base resolution mapping of RNA 2-O-methylated residues. Nucleic Acids Res 45, 1433–1441.

Isaac, C., Yang, Y. and Meier, U. T. (1998). Nopp140 functions as a molecular link between the nucleolus and the coiled bodies. J. Cell Biol. 142, 407–417.

Khan, M. S. and Maden, B. E. (1978). Conformation of methylated sequences in HeLa cell 18-S ribosomal RNA: nuclease S1 as a probe. Eur J Biochem 84, 241–50.

Kiss, T. (2004). Biogenesis of small nuclear RNPs. J Cell Sci 117, 5949–51.

Krogh, N., Jansson, M. D., Hafner, S. J., Tehler, D., Birkedal, U., Christensen-Dalsgaard, M., Lund, A. H. and Nielsen, H. (2016). Profiling of 2’-O-Me in human rRNA reveals a subset of fractionally modified positions and provides evidence for ribosome heterogeneity. Nucleic Acids Res 44, 7884–95.

Lafontaine, D. L. (2015). Noncoding RNAs in eukaryotic ribosome biogenesis and function. Nat Struct Mol Biol 22, 11–9.

Lemm, I., Girard, C., Kuhn, A. N., Watkins, N. J., Schneider, M., Bordonne, R. and Luhrmann, R. (2006). Ongoing U snRNP biogenesis is required for the integrity of Cajal bodies. Mol Biol Cell 17, 3221–31.

Lestrade, L. and Weber, M. J. (2006). snoRNA-LBME-db, a comprehensive database of human H/ACA and C/D box snoRNAs. Nucleic Acids Res 34, D158–62.

Maden, B. E. (1972). Effect of amino acid starvation on ribosome formation in HeLa cells. Ribosomal labelling patterns in cells deprived of different individual amino acids. Biochim Biophys Acta 281, 396–8.

Maden, B. E. (1986). Identification of the locations of the methyl groups in 18 S ribosomal RNA from Xenopus laevis and man. J Mol Biol 189, 681–99.

Maden, B. E. (1988). Locations of methyl groups in 28 S rRNA of Xenopus laevis and man. Clustering in the conserved core of molecule. J Mol Biol 201, 289–314.

Maden, B. E., Corbett, M. E., Heeney, P. A., Pugh, K. and Ajuh, P. M. (1995). Classical and novel approaches to the detection and localization of the numerous modified nucleotides in eukaryotic ribosomal RNA. Biochimie 77, 22–9.

Maden, B. E. and Salim, M. (1974). The methylated nucleotide sequences in HELA cell ribosomal RNA and its precursors. J Mol Biol 88, 133–52.

Mahmoudi, S., Henriksson, S., Weibrecht, I., Smith, S., Soderberg, O., Stromblad, S., Wiman, K. and Farnebo, M. (2010). WRAP53 is essential for Cajal body formation and for targeting the survival of motor neuron complex to Cajal bodies. PLoS Biol 8, e1000521.

Marcel, V., Catez, F. and Diaz, J. J. (2015). Ribosome heterogeneity in tumorigenesis: the rRNA point of view. Mol Cell Oncol 2, e983755.

Marcel, V., Ghayad, S. E., Belin, S., Therizols, G., Morel, A. P., Solano-Gonzalez, E., Vendrell, J. A., Hacot, S., Mertani, H. C., Albaret, M. A. et al. (2013). p53 acts as a safeguard of translational control by regulating fibrillarin and rRNA methylation in cancer. Cancer Cell 24, 318–30.

Meier, U. T. (2016). RNA modification in Cajal bodies. RNA Biol, 1–8.

Poole, A. R., Enwerem, II, Vicino, I. A., Coole, J. B., Smith, S. V. and Hebert, M. D. (2016). Identification of processing elements and interactors implicate SMN, coilin and the pseudogene-encoded coilp1 in telomerase and box C/D scaRNP biogenesis. RNA Biol 13, 955–972.

Raimer, A. C., Gray, K. M. and Matera, A. G. (2016). SMN - A chaperone for nuclear RNP social occasions? RNA Biol, 1–11.

Schattner, P., Brooks, A. N. and Lowe, T. M. (2005). The tRNAscan-SE, snoscan and snoGPS web servers for the detection of tRNAs and snoRNAs. Nucleic Acids Res 33, W686–9.

Schimmang, T., Tollervey, D., Kern, H., Frank, R. and Hurt, E. C. (1989). A yeast nucleolar protein related to mammalian fibrillarin is associated with small nucleolar RNA and is essential for viability. Embo J 8, 4015–24.

Shpargel, K. B. and Matera, A. G. (2005). Gemin proteins are required for efficient assembly of Sm-class ribonucleoproteins. Proc Natl Acad Sci U S A 102, 17372–7.

Stanek, D. (2016). Cajal bodies and snRNPs - friends with benefits. RNA Biol, 1–9.

Szewczak, L. B., DeGregorio, S. J., Strobel, S. A. and Steitz, J. A. (2002). Exclusive interaction of the 15.5 kD protein with the terminal box C/D motif of a methylation guide snoRNP. Chem Biol 9, 1095–107.

Trinkle-Mulcahy, L. and Sleeman, J. E. (2016). The Cajal body and the nucleolus: “In a relationship” or “It’s complicated”? RNA Biol, 1–13.

Tyc, K. and Steitz, J. A. (1989). U3, U8 and U13 comprise a new class of mammalian snRNPs localized in the cell nucleolus. EMBO J 8, 3113–9.

Tycowski, K. T., Aab, A. and Steitz, J. A. (2004). Guide RNAs with 5’ caps and novel box C/D snoRNA-like domains for modification of snRNAs in metazoa. Curr Biol 14, 1985–95.

Tycowski, K. T., Smith, C. M., Shu, M. D. and Steitz, J. A. (1996). A small nucleolar RNA requirement for site-specific ribose methylation of rRNA in Xenopus. Proc Natl Acad Sci U S A 93, 14480–5.

van Nues, R. W., Granneman, S., Kudla, G., Sloan, K. E., Chicken, M., Tollervey, D. and Watkins, N. J. (2011). Box C/D snoRNP catalysed methylation is aided by additional pre-rRNA base-pairing. EMBO J 30, 2420–30.

Wang, W., Nag, S., Zhang, X., Wang, M. H., Wang, H., Zhou, J. and Zhang, R. (2015). Ribosomal proteins and human diseases: pathogenesis, molecular mechanisms, and therapeutic implications. Med Res Rev 35, 225–85.

Watkins, N. J., Leverette, R. D., Xia, L., Andrews, M. T. and Maxwell, E. S. (1996). Elements essential for processing intronic U14 snoRNA are located at the termini of the mature snoRNA sequence and include conserved nucleotide boxes C and D. RNA 2, 118–33.

Yang, J. H., Zhang, X. C., Huang, Z. P., Zhou, H., Huang, M. B., Zhang, S., Chen, Y. Q. and Qu, L. (2006). snoSeeker: an advanced computational package for screening of guide and orphan snoRNA genes in the human genome. Nucleic Acids Res 34, 5112–23.

Yu, Y. T., Shu, M. D. and Steitz, J. A. (1998). Modifications of U2 snRNA are required for snRNP assembly and pre-mRNA splicing. Embo J 17, 5783–95.

Zhang, J., Zhang, F. and Zheng, X. (2010). Depletion of hCINAP by RNA interference causes defects in Cajal body formation, histone transcription, and cell viability. Cell Mol Life Sci 67, 1907–18.

Zhou, X., Liao, W. J., Liao, J. M., Liao, P. and Lu, H. (2015). Ribosomal proteins: functions beyond the ribosome. J Mol Cell Biol 7, 92–104.

